# Context dependent activation and repression of enhancers by Hunchback binding sites in *Drosophila* embryo

**DOI:** 10.1101/2020.10.21.348722

**Authors:** Stefano Ceolin, Monika Hanf, Max Schnepf, Ulrich Unnerstall, Christophe Jung, Ulrike Gaul

**Affiliations:** Gene Center and Department of Biochemistry, Center for Protein Science Munich (CIPSM), Ludwig-Maximilians-Universität München, Feodor-Lynen-Strasse 25, 81377 München, Germany; Faculty of Biology, Biozentrum, Ludwig-Maximilians-Universität München, 82152 Planegg-Martinsried, Germany

## Abstract

Hunchback (Hb) is considered a context-dependent transcription factor, able to activate or repress different enhancers during *Drosophila* embryo segmentation. The mechanism driving the contextdependent activity of Hb is however not well understood. Here we measure the activity of a large set of 20 synthetic enhancers that we design to elucidate the effect of Hb binding sites in *Drosophila* segmentation. We obtain quantitative data on the spatiotemporal dynamics of activity of all synthetic enhancers *in-vivo*, by using a quantitative and sensitive reporter system we recently developed. Our data reveal the dual role of Hb binding sites in segmentation enhancers: on the one hand, Hb act as a typical short range repressor by binding to its cognate sequences; on the other hand, we report a novel effect of a sequence containing multiple Hb binding sites, which is able to increase enhancer activity independently from Hb binding. This sequence, which contains multiple Poly-dA stretches, increases the activity of enhancers driven by different activators, possibly by disfavoring nucleosome occupancy.

**AUTHOR SUMMARY:** The control of gene expression is a fundamental process that allows cells to respond to external stimuli and take on various identities in complex organisms. Enhancers are DNA sequences that play a key role in this process. In the simplest model of an enhancer, small parts of its sequence can be specifically bound by proteins called transcription factors and the occupancy pattern of these proteins on the enhancer determines the expression level of a specific gene. In this research work we have studied enhancers in the context of the development of a fruit fly embryo. We have built synthetic enhancer sequences containing binding sites for a few specific factors and measured their activity in living embryos using fluorescence microscopy. Our results revealed that binding sites for a particular protein, Hunchback, are able to influence the activity of the enhancer even independently from Hunchback binding to them. This discovery might help to explain the complex effects that have been observed when studying Hunchback binding sites in natural enhancers.

## INTRODUCTION

Enhancers, or cis-regulatory elements (CREs), are DNA elements recognized and bound by multiple transcription factors (TFs), thus tuning transcriptional initiation of target genes. The spatiotemporal control held by enhancers is pivotal in the development of complex organisms, and sequence variations in these regulatory elements have been linked to a variety of human conditions (1, 2). The establishment of body segments along the antero-posterior (A-P) and dorso-ventral (D-V) axes during *Drosophila Melanogaster* (*D. Melanogaster*) development is controlled by a network of enhancers, and has longed served as a fundamental paradigm to study transcriptional regulation (3). However, even in this well studied paradigm, our understanding of how the enhancers function remains limited. For example, it is not yet possible to reconstitute entire enhancers from scratch by combining what are thought to be their key component parts (4).

Enhancers activity is ultimately determined by their sequence, which typically contains a complex architecture of clusters of binding sites for various TFs. Gene expression is a complex function of these sites’ occupancy, integrating the opposing effects of factors that either promote or inhibit transcription (5, 6). How this architecture of binding sites determines expression has been the subject of intensive investigation, with the ultimate goal to develop predictive models linking an enhancer’s sequence to its activity. Fractional occupancy models have been remarkably successful in predicting enhancer’s activity (7), despite their strong simplifying assumptions. The use of synthetic enhancers has proved useful in characterizing various properties of enhancer’s architecture. This allowed for example to elucidate the role of cooperativity among closely spaced binding sites for the maternal activator Bicoid (Bcd) (8, 9) or to implement a fully synthetic enhancer system to study how combination of activators and repressors generate precise expression patterns (10). However, failures in the attempt of reconstituting enhancers by combining transcription factor binding sites (4) and the difficulties in interpreting the results of some synthetic enhancers experiments (11) suggest that the simple binding preferences of activating/repressing factors are not the sole determinant of enhancer activity. Experimental hints (12-15) clearly indicate that a more complex level of regulation, involving the arrangement and spacing of binding sites as well as additional features of the DNA context sequence (the bases in between and surrounding the TF binding sites), also play a key role. Multiple biochemical mechanisms have been suggested to be involved in this process. Binding sites occupancy can be promoted by cooperativity (12), or hindered by competition with other factors (16) and by the presence of nucleosomes. Enhancers have been found to overlap with nucleosome depleted regions (6, 14, 15), thus pointing to a role of DNA accessibility as a prerequisite for the binding of some input TFs to their cognate sites. Nucleosome sequence preferences might also be important to control gene expression. For example, Poly-dA-dT sequences are considered to be the strongest nucleosome disfavoring motifs (17-19) and they have been found to influence the activity of promoters by reducing nucleosome occupancy in their vicinity (20).

TFs binding can also affect enhancer activity through chromatin mediated mechanisms. It has been known for a while that short and long range transcriptional repressors are able to increase nucleosome occupancy and induce distinct chromatin states either locally (over ∼100bp), or at the level of an entire locus (21, 22). In contrast, the pioneer transcription factor Zelda, which is a master regulator of the zygotic genome activation in *D. Melanogaster* development, is known to promote an open chromatin structure in the region surrounding its binding sites (23-25). Taken together these observations suggest a scenario in which TFs and nucleosomes dynamically influence each other, in a process that is ultimately guided by the enhancer sequence, and that determines enhancer activity.

Considering the complex biochemical processes underlying transcriptional regulation, it comes as no surprise that the effect of some TFs involved in *D. Melanogaster* segmentation is context dependent. One particularly relevant example is given by Hb, a key regulator of many gap (26) and segment polarity genes (16, 27) of *Drosophila* segmentation network which also has a human homolog central for human hematopoiesis (28). In the early stages of *Drosophila* development Hb is expressed, both zygotically and maternally, in the anterior half of the embryo and has been reported to act as an activator or a repressor on different enhancers, as for example the eve2+7 and eve3+7 enhancers (29, 30). This context dependent behavior has been observed even for enhancer active in the same cells, thus implying that some features of the enhancer sequence control Hb bifunctionality (30). Early reports postulated that Hb could switch from a repressive to an activating factor when bound to the enhancer in close proximity to Bicoid (31). Other studies have instead suggested that different binding modalities of Hb (e.g. as dimers vs monomers) could be a key determinant of its effect on gene expression (32). Even when Hb is clearly acting as a repressor (for example in setting the anterior boundary of activity of various gap or segment-polarity enhancers (Kruppel (Kr), Knirps (Kni), Giant (Gt) and others) its repressive activity can show different context dependent behavior. While for some of these enhancers Hb is sufficient for repression and works in a simple concentration dependent manner, for other enhancers Hb only creates a permissive environment for the action of additional repressive factors (13). A potential explanation is that Hb could be a short-range repressor, for which the arrangement and spacing of binding sites is critical in determining their effect on enhancer activity (33). Moreover, the ability of Hb to create a permissive environment for the action of other factors could be linked to its ability to recruit chromatin remodeler dMi-2, a component of the NURD chromatin remodeling and de-acetylation complex (34). A recent work questioned some of these observations and attributed Hb bifunctionality observed in Hb misexpression experiments (30), to Hb counter-repressing other transcriptional repressors (35). Despite this relatively large set of observations we haven’t yet reached a satisfactory understanding of the role of Hb binding site in segmentation enhancers and, in particular, how different features of enhancers sequence can coordinate the different behaviors of Hb.

Synthetic biology could offer the potential to clarify the role of Hb binding sites in enhancers by measuring the activity of synthetic sequences carrying different arrangements of binding sites for Hb combined with other activator TFs. However, to fully exploit the potential of synthetic enhancer constructs it is necessary to precisely track their activity in space and time with a system that is sensitive enough to measure both weak and strong enhancers, and to reliably detect subtle quantitative effects. This proves to be challenging using standard *in-situ* hybridization staining techniques because of their qualitative or semi-quantitative readout of mRNA concentration, which is due to the non-linear nature of enzymatic labeling. We recently developed an optimized version of the bright and fast-maturing fluorescent protein mNeonGreen to serve as a real-time, quantitative reporter of gene/enhancer expression (36). This approach employs a robust reconstruction algorithm to derive instantaneous and cumulative mRNA production rates from the dynamics of reporter fluorescence with high spatial and temporal resolution. This reporter system proved to be advantageous in terms of sensitivity and throughput compared to previous methods (36), and it is ideally suited to quantitatively study the activity of tens of synthetic constructs *in vivo*.

In this work, we obtain new insights about the role of Hb binding sites by measuring the spatiotemporal dynamics of 20 synthetic enhancer sequences combining binding sites for Hb with those of different activators. First, following a similar experimental strategy as introduced by Fakhouri et al. (21), we combined Hb binding sites with sites for two orthogonal activating transcription factors of the DV patterning system Twist (Twi) and Dorsal (Dl), which are homogenously expressed in the ventral side of the embryo. This setting allows us to study the impact on the enhancer’s activity of binding sites spacing and orientation. Importantly, since the Hb protein is only expressed in the anterior part of the embryo, we are able to simultaneously monitor the enhancer activity both in the presence or absence of the Hb protein in a single embryo. In a second set of experiments, we exchanged the Twi-DI binding sites with binding sites for a different TF, the anterior activator Bcd, and found similar effects of the Hb binding sites on expression. Our results reveal a dual role of Hb binding sites in shaping segmentation enhancers activity: on the one hand, Hb can act as a typical short range repressor by binding to its cognate sequences; on the other hand, we report a novel effect of a sequence containing multiple Hb binding sites, which enhances expression of both Twi-Dl and Bcd driven enhancers independently from Hb binding. As this sequence contains multiple Poly-dA stretches, it may promote a permissive environment for the enhancer activity by disfavoring nucleosome occupancy, leading to enhanced enhancer activity. Overall, we propose that the net effect on expression of including Hb binding sites in a segmentation enhancer depends on a balance between the activating effect of the binding sites themselves and the repression of Hb binding to those sites. However, for all the enhancers tested in this work we observed a boost to enhancer activity from the insertion of Hb binding sites. Moreover, the distance dependencies of Hb repression and Hb binding sites activation are different, thus creating a strong non-linear behavior of enhancer’s activity as a function of enhancer’s architecture.

## RESULTS

### Spatiotemporal characterization of enhancer activity

We started by measuring the activity of a well-established 57bp-long synthetic construct derived from the *snail* proximal enhancer, and containing two binding sites for each of the D-V activators Twist and Dl (2Twi-2Dl; **Fig. 1A**) (37). The activity of this sequence has been characterized using *in-situ* hybridization staining and exhibits homogenous activity in the embryo ventral side (21, 37).

**Figure 1.**
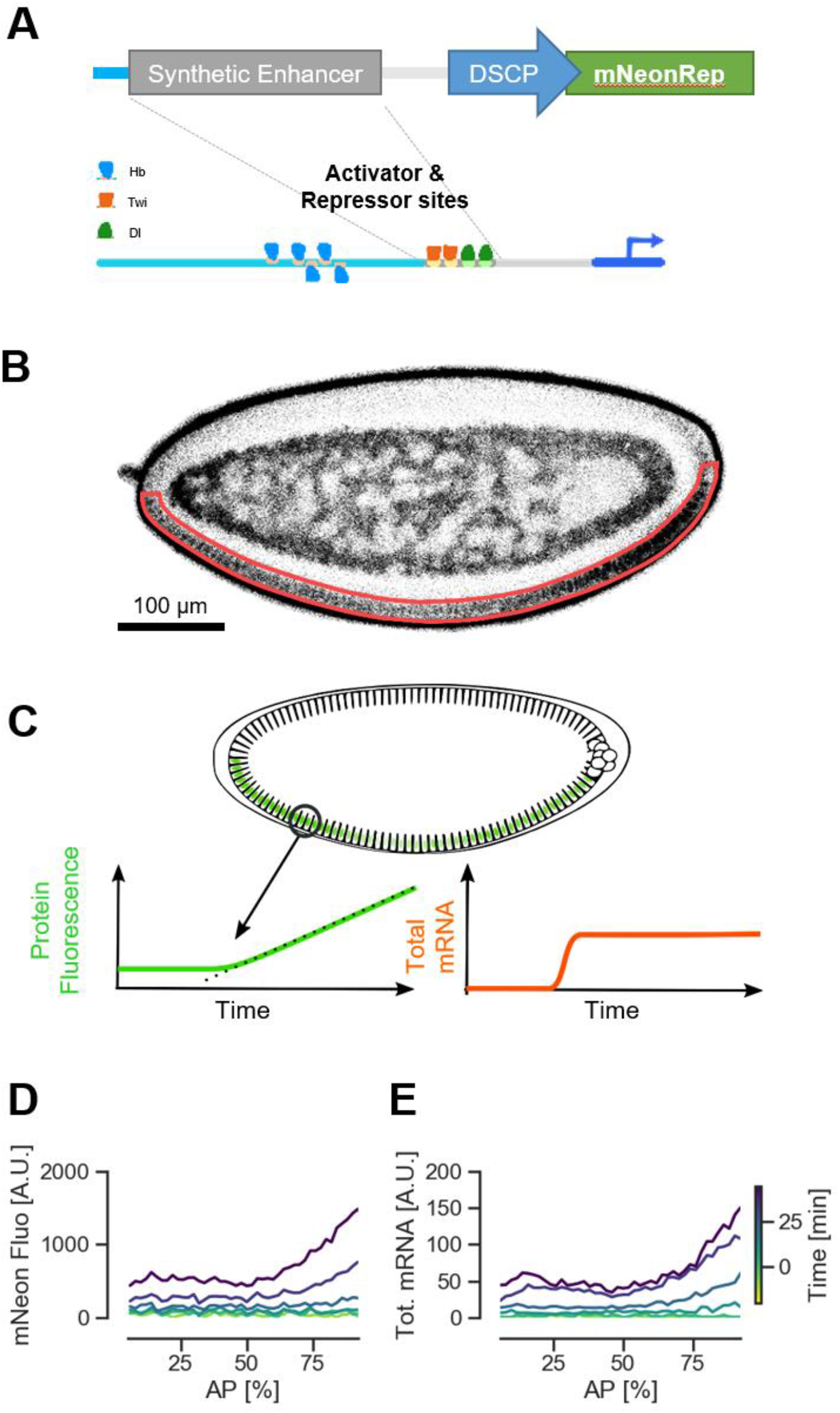
Quantifying synthetic enhancers activity with the mNeonRep reporter. **A)** Synthetic regulatory sequences containing binding sites for selected transcription factors are cloned upstream a DSCP promoter controlling the expression of the mNeon reporter. **B**) Representative confocal slice of a living embryo, laid by homozygous transgenic parents carrying the reporter construct (Original-2Twi-2Dl-mNeonRep), showing mNeon fluorescence in grayscale. The fluorescence of the mNeon reporter can be detected in the nuclei in cortical region of the embryo, on the ventral side (highlighted in red in the figure). **C**) Sketch illustrating the use of a fluorescent protein as a transcriptional reporter. The time course of protein fluorescence carries information about the underlying dynamics of mRNA production, which can be recovered with a reconstruction algorithm. **D**) mNeon fluorescence patterns along the AP axis of embryo, on the ventral side. mNeon fluorescence has been quantified from three confocal slices and averaged in bins corresponding to 2% of the embryo length. Each line corresponds to a different time of embryo development. **E**) Patterns of cumulative mRNA production reconstructed from the time course of mNeon fluorescence.

Since for small enhancers, the surrounding sequence could play an important role, at first we included upstream of the enhancer ∼300bp of the flanking background sequence from the original reporter plasmid used in Fakhouri et al. (21), which we found to carry a substantial number of binding sites for TFs of the segmentation network, in particular for Hb (**Fig. 1A** and **Supplementary Fig. 1**). As for all enhancer constructs presented in this study, to detect subtle quantitative effects in the activity of different constructs we used a new reporter system that we have recently developed (36). This reporter is particularly well suited for studying the weak activity of short synthetic enhancers in *D. Melanogaster* blastoderm embryo. Briefly, the mNeon reporter system is based on the expression of an optimized reporter fluorescent protein and on a data analysis pipeline that infers the information about mRNA levels by analyzing the time course of protein fluorescence with a model of ordinary differential equations (ODEs) (38). As reported by other studies (21), we observe that the 2Twi-2Dl enhancer drives relatively weak and homogenous expression on the ventral side of the embryo (**Fig. 1B, D** and **E**). However, our data also show a somewhat stronger activity towards the embryo posterior, which may not have been evident using *in situ* staining.

### The effect of Hb binding sites in Twist-Dorsal driven enhancers

The effect of Hb binding sites has been mainly investigated in the context of complex natural enhancers like the *eve* enhancers (35, 39). To analyze their effect systematically, we started by designing synthetic enhancer constructs combining Hb binding sites with the 2Twi-2Dl enhancer. Moreover, to work in a clean setting without binding sites for any other segmentation TFs, we introduced in all our constructs right upstream the enhancer a different 340bp neutral flanking sequence. This spacer sequence does not contain any predicted binding sites for any segmentation TFs, as checked using PySite (available on *Github*: https://github.com/Reutern/PySite) a *Python* script we developed which uses positional weight matrices (PWMs) that we determined in a previous work (40) (**Supplementary Fig. 2**). The spacer sequence has also been proven not to drive any expression *in vivo* (41). In addition to a Twi and Dl construct having the neutral background sequence (2Twi-2Dl; **Fig. 2B**), we generated 4 additional ennhancers by introducing a sequence containing 3 functional binding sites for Hb (31). In particular we inserted the sites just upstream the 2Twi-2Dl enhancer (3Hb-2Twi-2Dl; **Fig. 2C**), or at increasing distance: 70bp (3Hb-70-2Twi-2Dl; **Fig. 2D**), 150bp (3Hb-150-2Twi-2Dl; **Fig. 2E**) and 250bp (3Hb-250-2Twi-2Dl ; **Fig. 2F**) away from the activator sites. Surprisingly, when exchanging the flanking sequence used in older studies with the neutral background, expression was substantially reduced to a barely detectable level (**Fig. 2A, B**). In addition, introducing 3 Hb sites in this new setting substantially increased enhancer activity in the entire embryo (**Fig. 2C, J**). Interestingly, the strong activating effect of Hb binding is not localized to anterior half of the embryo where Hb is present and, therefore, cannot be attributed to Hb binding. Moreover, the expression pattern of the 3Hb-2Twi-2Dl enhancer is not homogenous along the AP axis. In particular, the region where Hb is expressed (represented with a gray shading in **Fig. 2J**) corresponds to a region of relatively weaker expression, around ∼3 times weaker compared to the posterior region. However, even in this region, the overall balance of inserting 3Hb binding sites in the enhancer still results in an increased activity compared to the 2Twi-2Dl enhancer (blue curve in **Fig. 2J**). This could be attributed to the combination of an activating effect from the presence of Hb binding sites and of the repressive effect of Hb binding to them.

**Figure 2.**
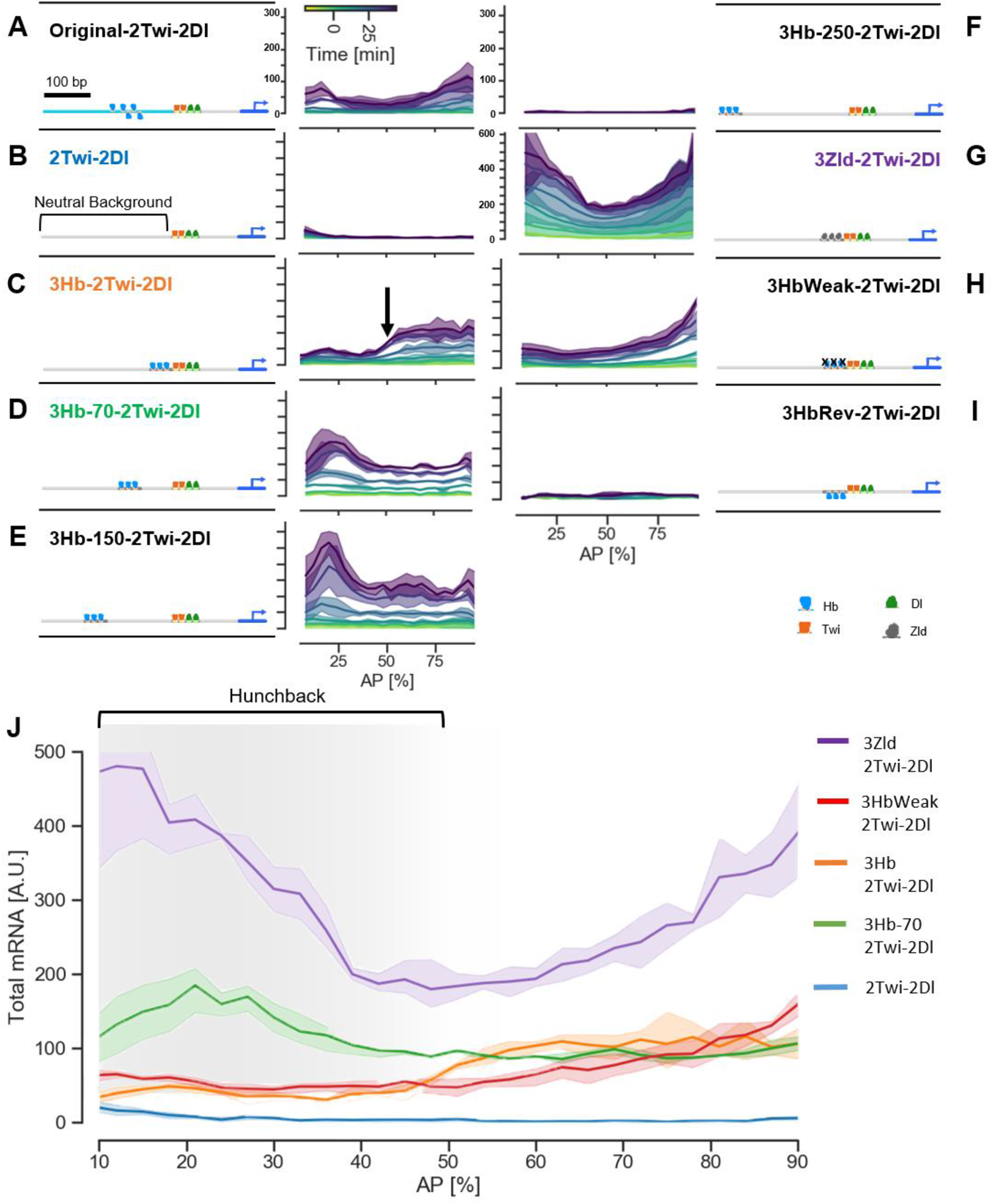
Overview of Twist and Dorsal driven synthetic enhancers expression patterns. **A-I)** Spatial-temporal dynamics of the activity of Twi and Dl driven synthetic enhancers. Solid lines represent the average cumulative mRNA production from 2 or 3 embryos grouped in bins corresponding to 4% of embryo length. The shaded areas represent ±1σ confidence intervals. All data represent the reporter expression in the ventral side of the embryo. **J**) Comparison of total cumulative mRNA production driven by a subset of enhancers. The gray shading represents the area of the embryo in which Hb is expressed. The arrows highlight the activating effect of the 3Hb sequence, and the relative repression due to Hb binding in the embryo anterior.

The expression pattern significantly varies when the distance between the 3 Hb and the activator sites is increased, with important differences between the anterior and posterior half of the embryo. In the posterior region, the activating effect of Hb binding sites remains constant by increasing the distance to 70bp or 150bp (**Fig. 2D-E**), while for a larger distance (250bp) any activating effect is lost and the expression returns to the baseline level (**Fig. 2F**). The anterior part of the embryo shows a very different behavior. The relative repression that we attribute to Hb binding is lost when the 3 Hb and Twi-Dl sites are separated by 70bp, the expression becoming even stronger than in the posterior. Anterior activity increases further when the separation is increased to 150bp. Finally, also in this part of the embryo, the expression returns to baseline level when the distance is further increased to 250bp. These observations in the embryo anterior, together with the overlap of the Hb expression domain with the region of relative repression for the 3Hb-2Twi-2Dl, point to Hb acting as a typical short-range repressor while the sequence of Hb bindings sites fosters enhancer activity.

To further investigate whether the relative repression of the 3Hb-2Twi-2Dl enhancer in the embryo anterior is indeed due to Hb binding, we introduced single point mutations in each of the 3Hb binding sites. In particular we mutated the core of each Hb binding site switching A->T, thus leading to a 10 folds decrease in the predicted binding site strength, without affecting the enhancer GC content. The activity of this mutated construct doesn’t show any clear drop in correspondence of the Hb expression domain (3HbWeak-2Twi-2Dl; **Fig. 2H, J**). Therefore, point mutations in the Hb motif inhibit Hb binding and consequent repression.

The interesting observation that a sequence containing 3Hb binding sites increases enhancer activity without binding of Hb could indicate that it carries DNA features promoting a permissive state of the enhancer. This could be achieved, for example, by influencing the enhancer accessibility. In this respect, an interesting observation is that Hb binding preferences coincide with a stretch of As(40) that constitutes a poly-dA-dT sequence. Poly-dA-dT sequences are considered to be the strongest nucleosome disfavoring motifs (17, 18) and they have been found to influence the activity of promoters by reducing nucleosome occupancy in their vicinity To gather additional hints in this direction, we tested if the enhancer sequence with only Twi and Dl binding sites is in a relatively inaccessible state, or if it is already highly accessible. To this end, we inserted 3 binding sites for the pioneer TF Zelda, which is known to increase enhancer accessibility. Among the synthetic enhancers studied in this work, this construct drove the highest activity along the whole AP axis (**Fig. 2G)**, suggesting that indeed the *2Twi-2Dl* enhancer weak activity is affected by a low accessibility. Moreover, to investigate if the 3Hb sequence has the potential to significantly influence nucleosome occupancy, we computationnaly predicted nucleosome occupancy based on *in vitro* nucleosome sequence preferences, by using a model developed by Kaplan et al. (17). This model predicts a substantially different nucleosome occupancy landscape among the synthetic enhancers we studied (**Supplementary Figs. 1, 2 and 3**). In particular, the 2Twi-2Dl enhancer containing the original flanking sequence has a low nucleosome occupancy (**Supplementary Fig. 1B**). In contrast, the 2Twi-2Dl enhancer with a clean background sequence, which has a very low activity, has a high predicted nucleosome occupancy (**Supplementary Fig. 2B**). The insertion of the 3Hb sequence into this enhancer, substantially reduces nucleosome occupancy (**Supplementary Fig. 3B**), as expected given that Poly (dA) tracts have the lowest average nucleosome occupancy (17). To further support these observations we used a fluorescence anisotropy assay we recently developed (42) to measure the *in vitro* nucleosome binding energy of three 150bp long DNA sequences, encompassing either the 2Twi-2Dl, 3Hb-2Twi-2Dl or 3HbWeak-2Twi-2Dl enhancers and part of their flanking sequence. Even though the differences of nucleosome binding energies among these enhancer sequences are not statistically significant, we could observe that nucleosome binding energy increases with enhancer activity (see **Supplementary Fig**.**6**).

Finally, we also explored the effect of the orientation of the Hb binding sites. It is generally accepted that enhancer activity is not sensitive to orientation. Surprisingly, we observed a major impact of the orientation of Hb binding sites: reversing the orientation of the sequence containing 3Hb sites substantially reduced expression in the entire embryo (3HbRev-2Twi-2Dl; **Fig. 2I**). Such an effect of orientation on expression, although weaker, has already been reported for Poly-dA tracts in yeast promoters (43).

### The effect of Hb binding sites in Bicoid driven enhancers

To test if the observed effect of Hb binding sites is consistent in different enhancers containing binding sites for other activators besides Twi and Dl, we designed a second group of enhancers. We chose to combine the same set of Hb binding sites with sites for the Bcd activator. This design is more directly relevant for the understanding of native enhancers, since Hb and Bcd often regulate the same enhancers. Moreover, Hb bifunctionality has been originally reported in a similar setting, leading to postulate a synergistic effect between these two factors (11, 31). We expect the activity of these enhancers to be driven by Bcd binding. Unfortunately, since both Hb and Bcd are localized in the embryo anterior, in the same set of experiments it is not possible to observe the effect of unoccupied Hb binding sites on Bcd dependent enhancer activation without perturbing the enhancer sequence to weaken the binding sites.

Similarly to the synthetic constructs driven by the Twi and Dl activators, we inserted the sequence containing 3 Hb binding sites at increasing distances from a sequence carrying 3 binding sites for Bcd (**Fig. 3**). The enhancer with only 3 bcd binding sites (Bcd3) drives expression in the anterior most region of the embryo (Bcd3; **Fig. 3A**), as previously reported by other studies (44). Introducing 3 Hb binding sites just upstream the 3 Bcd sites slightly but significantly (p = 0.0048, two-tailed Welch’s t-test) increased enhancer activity (3Hb-Bcd3; **Fig. 3B, E**). This is in agreement with our previous observation that, at short range, the balance between the activating effect of Hb binding sites and the repressing effect of Hb binding to them is shifted towards activation.

**Figure 3.**
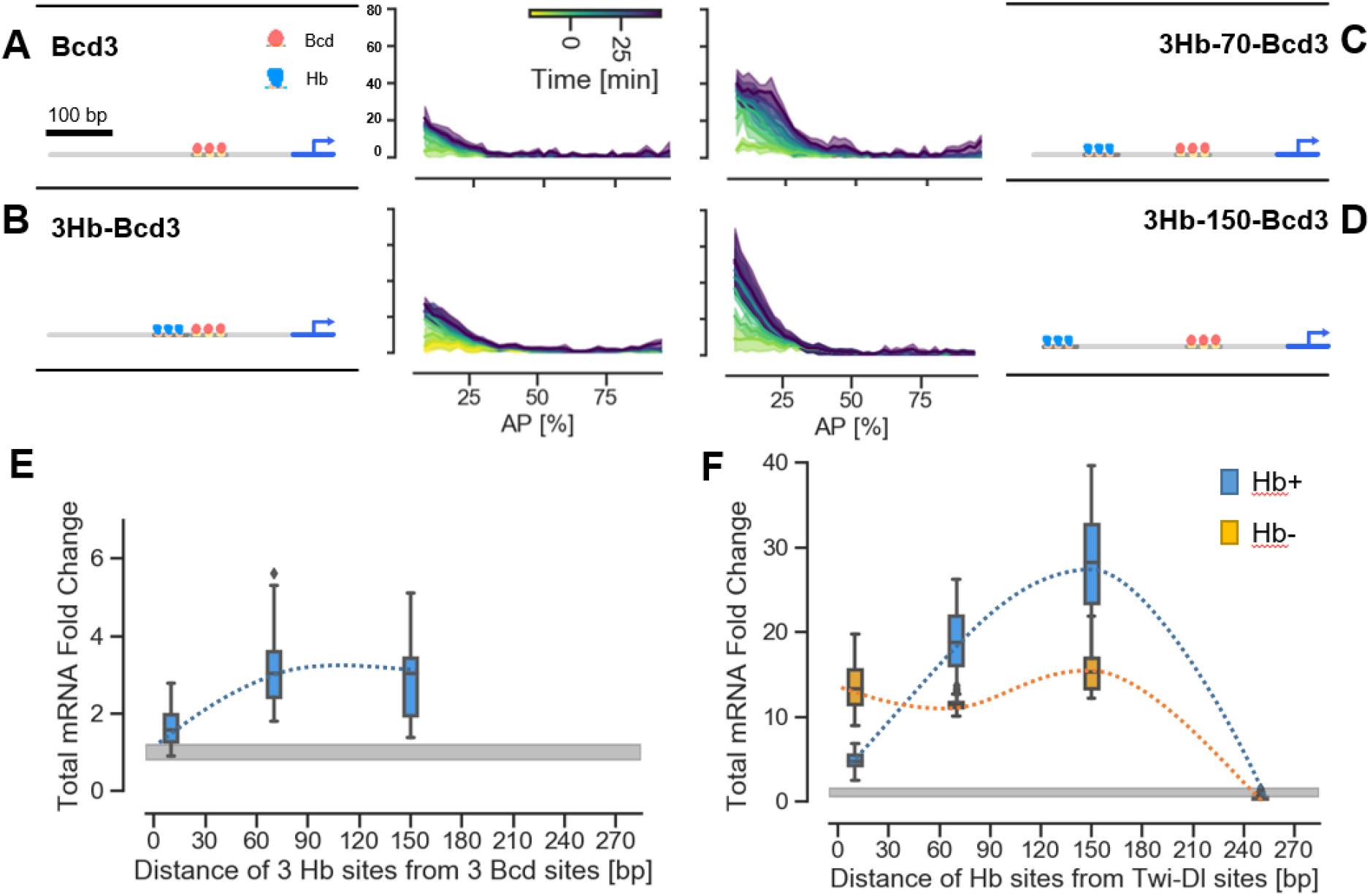
Hb binding sites similarly affect Bcd and Twi-Dl activity. Spatial-temporal dynamics of the activity of **A**) Bcd3 **B**) 3Hb-Bcd3 **C**) 3Hb-70-Bcd3 and **D**) 3Hb-150-Bcd3 synthetic enhancers. Solid lines represent the average cumulative mRNA production from 2 or 3 embryos grouped in bins corresponding to 4% of embryo length. The shaded areas represent ±1σ confidence intervals. All data represent the reporter expression in the ventral side of the embryo. **E**) Fold change of the total cumulative mRNA production in the embryo anterior for the enhancers with 3 Hb binding sites at various distances from the 3 Bcd binding sites, compared to the Bcd3 enhancer. Fold change has been computed independently in bins corresponding to 2% of the embryo length and data have been then pooled together in the area 10-30%AP. The gray area illustrates the variability in the total cumulative mRNA production level for the Bcd3 enhancer and represent and interval of ±1σ **F**) Fold change of the total cumulative mRNA production in the embryo anterior (Hb+) and posterior (Hb-) for the enhancers with 3 Hb binding sites at various distances from the 2Twi and 2Dl binding sites, compared to the 2Twi-2Dl enhancer. Fold change has been computed independently in bins corresponding to 2% of the embryo length and data have been then pooled together in the area 10-30%AP and 70-90%AP. The gray area illustrates the variability in the total cumulative mRNA production level for the 2Twi-2Dl enhancer and represent and interval of ±1σ.

One previous study investigated the activity of the Bcd3 and the 3Hb-Bcd3 enhancers (31), reporting an expansion towards the embryo posterior of the expression domain of the Bcd3 enhancer, upon inserting three binding sites for Hb. Importantly, an expansion of the expression domain with a constant peak activity supports the idea that the enhancer reacts to a different concentration threshold of its activator Bcd. Moreover, a threshold change following the inclusion of 3Hb sites in the enhancer could indicate a direct cooperative interaction between Hb and Bcd. In order to compare our results with this previous report (31) we checked wether the insertion of the 3Hb sequence caused an expansion of the expression domain driven by the enhancer. In our data, including 3Hb sites increases expression homogeneously and doesn’t substantially alter the shape of the expression domain and therefore its boundary (**Supplementary Fig. 4A, B**). This difference compared to older studies might be due to artefacts introduced by *in situ* stainings which offered a high sensitivity, potentially higher than our reporter, but had a non-linear response that could cause saturation.

Similarly to what we observed for the Twi-Dl driven enhancers, activity increased substantially when we introduced a 70 (3Hb-70-Bcd3; **Fig. 3C**) or 150bp (3Hb-150-Bcd3; **Fig. 3D**) long neutral spacer sequence between the Hb and Bcd sites. To characterize this distance dependence in both groups of enhancers, we plotted the fold change of the total amount of mRNA produced by each enhancer as function of the distance between Hb and Twi-Dl or Bcd sites (**Fig. 3E**). For the Bcd driven enhancers we explored a window spanning from 10% to 30% AP, and calculated the fold change in bins corresponding of 2% of the embryo length. Similarly, for the Twi-Dl driven enhancers we considered two regions, one from 10% to 30% of the AP axis and the other from 70% to 90% of the AP axis, corresponding to the presence or absence of Hb, respectively (**Fig. 3F**). Interestingly, in the anterior part where the Hb protein is present the fold change of enhancer’s activity induced by the presence of Hb binding sites shows a similar trend for both Bcd or Twi-Dl driven enhancers, increasing in both cases when a spacer of 70bp or 150bp is introduced although with a different overall impact. Thus, the effect on expression of the different settings of the Hb binding sites are in overall independent from the tested activator.

To investigate if inserting the 3Hb sequence only affects the enhancer or can also directly influence the activity of the promoter, we inspected the activity of the Bcd3 and 3Hb-150-Bcd3 enhancers, and we created two new constructs in which we varied the enhancer-promoter distance to test the effect of this second distance parameter. Enhancer-promoter distance can significantly influence the expression level driven by an enhancer (36). However, if the 3Hb sequence does not directly influence the promoter, we expect that it will influence enhancer activity of the same relative amount for different enhancer-promoter distances. This is indeed what we observed: whereas the absolute amount of mRNA production is increased when the enhancer is closer to the promoter (compare **Supplementary Fig. 5A, B** and **Fig. 3A, D**), the fold change of activity due to the insertion of the 3Hb sequence at 150bp from the Bcd3 sites is the same in both cases (**Supplementary Fig. 5C**).

Using the same setting, we also explored the effect of changing the number, strength and orientation of Hb binding sites on activity. Removing one Hb binding site from the 3Hb-150-Bcd3 enhancer only slightly reduced enhancer’s activity, which remained sustained (**Fig 4A, B** and **G**). However, removing a second Hb binding site reduced the activity to a level compatible with the baseline level of the Bcd3 enhancer (**Fig. 4C, G**). Single point mutations in all three Hb binding sites further increased the activity, revealing that even at 150bp away Hb binding may still have some residual repressing effect (**Fig. 4D**). Reversing the orientation of 3Hb binding sites proved again to have a severe impact on expression, although less pronounced than in the case of the Twi-Dl activator. At 150bp from the activator sites (**Fig. 4E)**, the reversed 3Hb sequence was still able to increase expression compared to the baseline level, but the activity was only half of that obtained in the forward orientation (**Fig. 4B)**. Similarly, when we inverted the Hb sites just upstream the Bcd binding sites (**Fig. 4G**), expression was also reduced and turned out to be comparable to that of the Bcd3 sequence alone.

**Figure 4.**
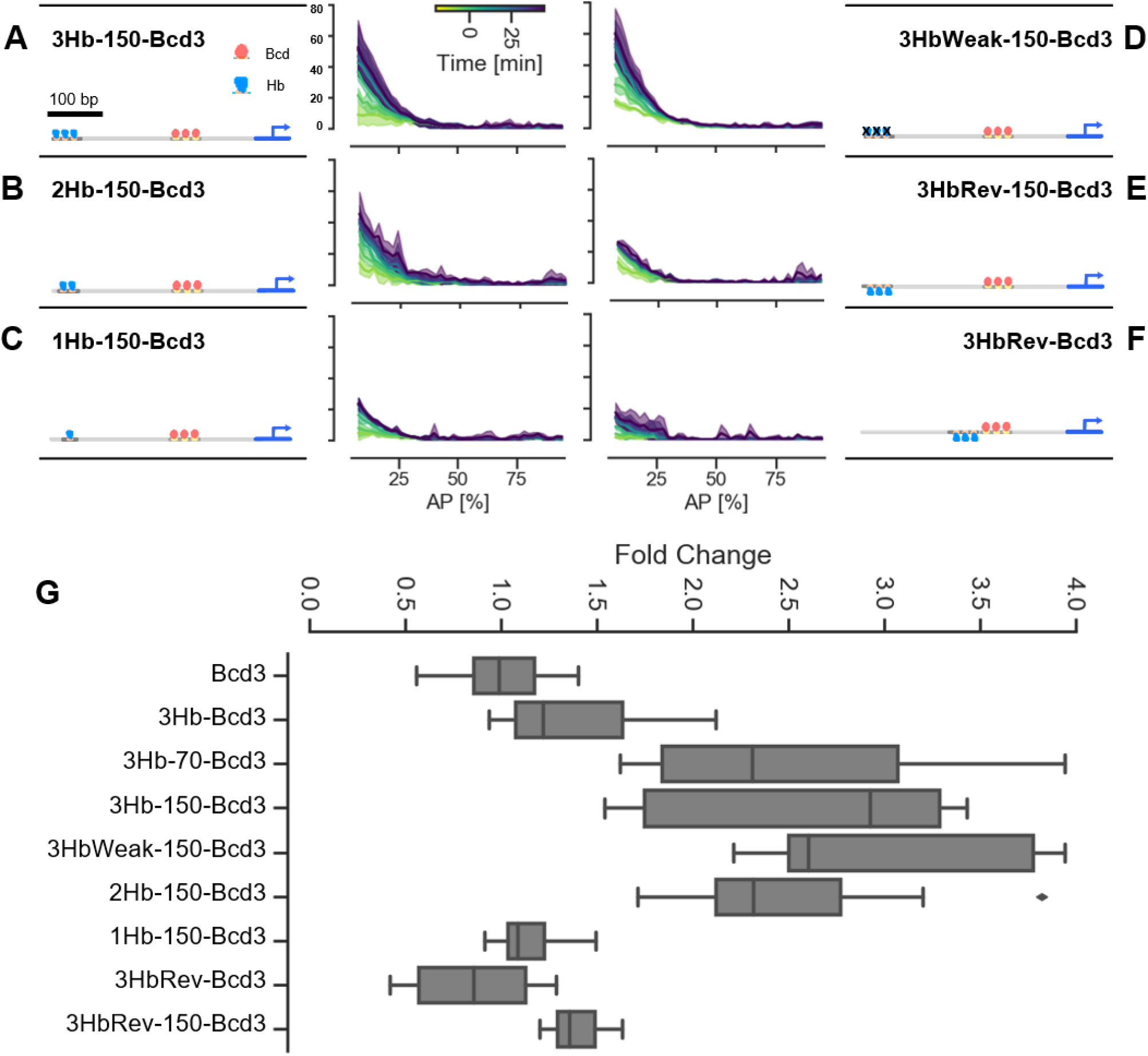
Effect of stoichiometry and orientation of Hb binding sites on Bicoid dependent activity. Spatial-temporal dynamics of the activity of **A**-**C**) synthetic enhancers containing 3, 2 or 1 binding sites for Hb at 150bp from the 3 Bcd sites. **D**) Activity of a synthetic enhancer containing 3 weaker binding sites for Hb obtained with point mutations of the consensus binding motif. **E-F**) synthetic enhancers containing 3 binding sites for Hb in reverse orientation either right upstream or at 150bp from the 3 Bcd sites. Solid lines represent the average cumulative mRNA production from 2 or 3 embryos grouped in bins corresponding to 4% of embryo length. The shaded areas represent ±1σ confidence intervals. All data represent the reporter expression in the ventral side of the embryo. **G**) Fold change of the total cumulative mRNA production in the embryo anterior for all Bcd-driven synthetic enhancers. Fold change has been computed independently in bins corresponding to 2% of the embryo length and data have been then pooled together in the area 10-30%AP.

Overall, it is remarkable that we found consistent effects when including a sequence containing 3 binding sites for Hb into synthetic enhancers based on different activators. In particular, in the anterior part of the embryo the inclusion in the enhancer of multiple Hb binding sites have in overall the same activating effect on the Twi-Dl and Bcd activators. We propose that this net activating input stems from two opposing mechanisms: short-range repression due to Hb binding to its cognate sites, coupled to the activating effect of Hb binding sites, which may be due to an increase in enhancer accessibility.

### Synthetic enhancers dynamics

Finally, we took advantage of the time resolution of the mNeon reporter system to investigate the temporal dynamics of our synthetic enhancers. To better highlight the time at which each enhancer is active, we used the instantaneous rate of mRNA production rather than the cumulative mRNA production. Our data clearly show that all Twi and Dl driven enhancers have a similar dynamics in the embryo anterior (**Fig. 5A**) and posterior (**Fig. 5B**). Moreover, they are activated at later time compared with the Bcd driven synthetic enhancers (**Fig. 5C**). The observed dynamics are consistent with the TFs concentration dynamics reported in prevuous studies: while Twi and Dl concentration rises through early embryo development and reaches a peak at the end of n.c. 14 (45, 46), Bcd concentration instead declines (45) and its activity is further decreased by Bcd sumoylation (47). Importantly, we couldn’t observe any consistent effect on the temporal dynamics caused by the insertion of Hb binding sites in any of the constructs under study. The only construct that shows a marked difference in its dynamics is the 3Zld-2Twi2Dl enhancer which is active much earlier than all other Twi and Dl driven enhancers (**Fig. 5A, B**). This observation also agrees with a study in which Zld binding sites were added to the snail enhancer, substantially accelerating enhancer activation (48).

**Figure 5.**
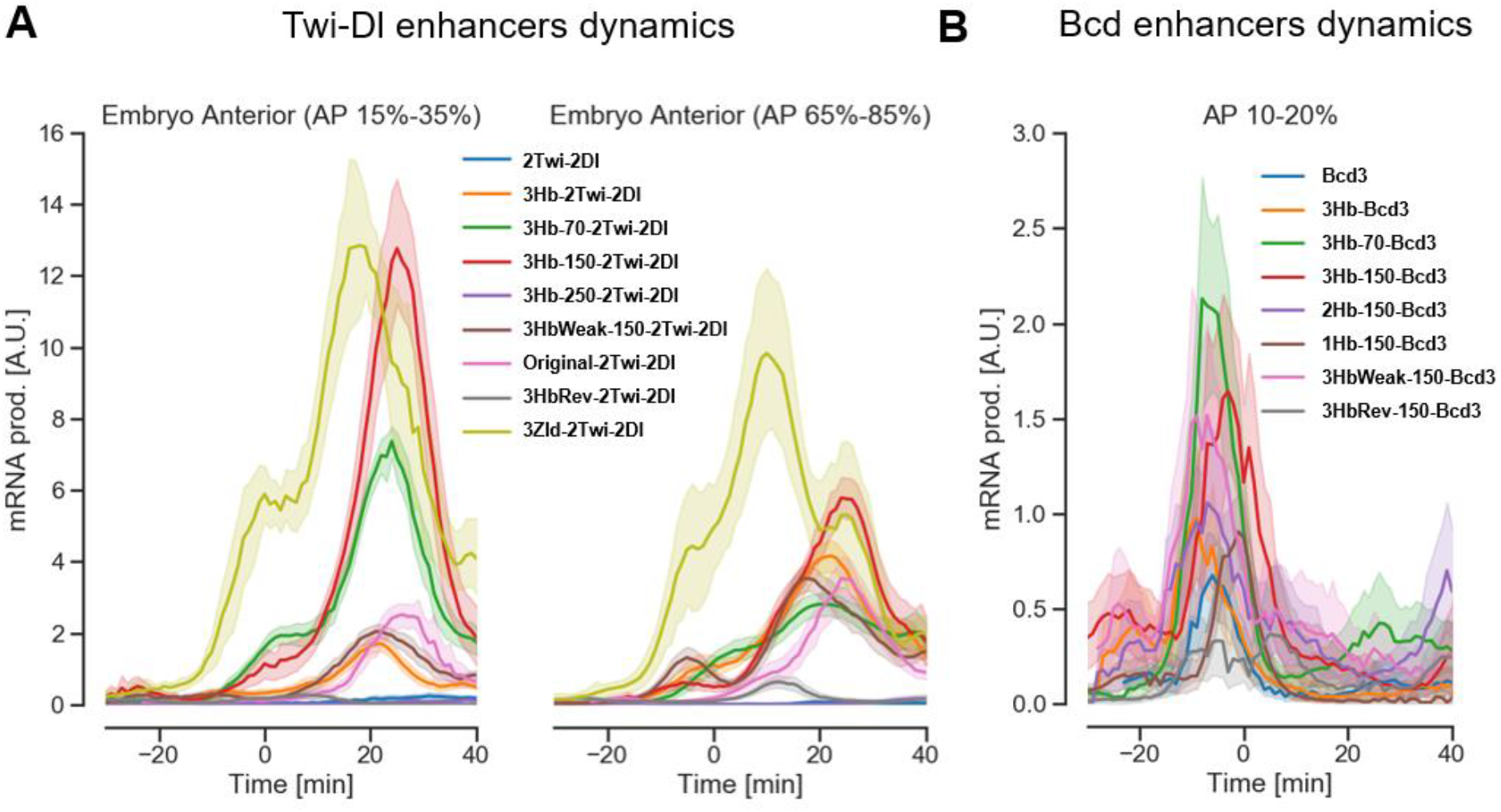
Temporal dynamics of synthetic enhancers expression. Average rate of instantaneous mRNA production reconstructed from the time course of protein fluorescence for all the enhancers in this study. The shaded areas represent ±1σ confidence intervals. **A**) temporal dynamics of enhancer activity for the Twi-Dl driven enhancers in the embryo anterior and posterior, respectively. **B**) temporal dynamics of Bcd driven enhancers in the embryo anterior.

## DISCUSSION

Interpreting the regulatory code of transcriptional enhancers requires an understanding of how multiple transcription factors and the chromatin mutually interact to modulate gene expression. This process is ultimately orchestrated by the DNA sequence of enhancers. Specifically, it is driven by the architecture of binding sites they contain and by the context sequence which can influence their activity. In this work we focused on Hb, a context dependent TF active in *D. Melanogaster* embryo segmentation. We quantitatively measured the transcriptional activity of 20 synthetic enhancers combining Hb binding sites with sites for either the dorso-ventral activators Twi and Dl, or the anterior activator Bcd. We found that (i) a sequence containing 3 Hb binding sites is able, independently from the presence of the Hb protein, to create a permissive environment for enhancer activity, strengthening the activity of other activators; (ii) when Hb binding sites are at close distance from the activator sites, Hb binding causes relative repression of enhancers activity, however, the overall balance of inserting the 3Hb sequence still leans towards activation; (iii) the 3Hb sequence acts consistently in both Twi and Dl or Bcd driven enhancers and, (iv) it influences enhancer activity levels without altering their temporal dynamics.

The expression patterns of synthetic and reconstituted enhancers have often been difficult to interpret (4, 11), although with some notable exceptions (21). In this work we show that using a quantitative and sensitive reporter to measure enhancer activity helps making further progresses. In fact, most of the differences we observed (in particular in the case of Bcd driven enhancers) are subtle quantitative effects on the expression of relatively weak enhancers. These effects may not have been resolved without the use of a reporter of enhancer activity that is both quantitative and sensitive, which is therefore pivotal in such a study. Fluorescent *in situ* hybridization stainings provide high sensitivity, but can only offer a qualitative or semi-quantitative readout of mRNA concentration, because of the non-linear nature of enzymatic labeling. More recently, new protocols based on advanced imaging techniques have been used to obtain quantitative data on mRNA expression, possibly even with higher sensitivity compared to the reporter used in this study. However, these methods are laborious, which can have an impact on experimental throughput.

Our findings also confirm that by using synthetic enhancers, great care must be taken in the selection of flanking and background sequences. We feel that flanking sequences in particular may have been overlooked in older studies. While their impact is probably limited when studying long native enhancers, flanking can strongly influence enhancer activity of shorter enhancers. The importance of flanking sequences of short enhancers is well illustrated by the comparison of the expression of the 2Twi-2Dl and 2Twi-2Dl-Original enhancers (see **Fig. 2 A** and **B**). The 2Twi-2Dl-Original enhancer included 300bp of the sequence flanking the 2Twi-2Dl enhancer in the reporter construct used by previous studies. This sequence contains multiple binding sites for Hb and increases enhancer activity. Importantly, this sequence is part of a plasmid backbone that has been largely used to generate reporter constructs to study enhancer expression (49).

Several studies identified Hb as a bifunctional TF: generally acting as a transcriptional repressor, but able to switch to an activator in specific contexts. These studies however didn’t reach an agreement regarding the mechanism driving this behavior. One study postulated that Hb bifunctionality might be driven by the formation of Hb dimers (32), similarly to what is known for other TFs. Others suggested that Hb bifunctionality would require specific protein-protein interactions with additional factors, for example when Hb binds to the enhancer in close proximity to a binding site for Bcd (31). Our data allow us to investigate more systematically this landscape, providing a more complete picture of the role of Hb binding sites in segmentation enhancers.

In particular, we report a novel effect of Hb binding sites which are, independently from Hb binding, able to influence enhancer activity. Specifically, we observed that a sequence containing 3Hb binding sites can substantially increase expression of enhancers in the embryo posterior, where Hb is only transiently expressed in a small domain and only late in blastoderm development. Moreover, the 3Hb sequence is not able to drive expression by itself (31), but it rather boosts the activity of other transcriptional activators. This is further confirmed by the observation that the activity of the synthetic enhancers studied here is always limited to the expression domains of the activating factors driving their expression.

An appealing explanation for the direct effect of the 3Hb sequence is that it could directly influence nucleosome occupancy, thus promoting the activity of transcriptional activators through an increase of DNA-accessibility. This model is supported by the following points: (i) the 3Hb sequence has the potential to significantly influence nucleosome occupancy. The predicted nucleosome occupancy of our synthetic enhancers is substantially reduced when the 3Hb sequence is included (**Supplementary Figs. 1, 2, 3**). (ii) The *in vitro* nucleosome binding energy of three 150bp long DNA sequences, encompassing either the 2Twi-2Dl, 3Hb-2Twi-2Dl or 3HbWeak-2Twi-2Dl enhancers (and part of their flanking sequence) correlate to some extent with enhancer activity (**Supplementary Fig. 6**). (iii) The distance dependence of the activating effect of the 3Hb sequence is compatible with what has been reported for the influence of Poly(dA:dT) tracts in promoters. The 3Hb sequence is able to influence expression even when positioned at 70 or 150bp away from the activator sites; it becomes ineffective only when the spacing is increased to 250bp. The observed distance dependence corresponds with that observed for the influence of Poly(dA:dT) tracts in the activity of yeast promoters (20), an effect that has been shown to be mediated by changes in nucleosome occupancy, and also to be sensitive to sequence orientation (43). Therefore, this behavior further supports the idea of Hb binding sites influencing enhancer activity by altering the enhancer accessibility. The effect of Poly(dA:dT) tracts has been already studied in the context of enhancers, in particular regarding their effect as flanking sequences of TF binding sites (50). In fact, Poly(dA:dT) can influence TF binding by their influence on DNA-shape. However, this effect is only observed if the Poly(dA:dT) tract is immediately adjacent to the binding sites, which is not the case in our synthetic constructs.

Our analysis of synthetic enhancers also confirmed the repressive activity of Hb, which we characterize as a short range repressor (**Fig. 2 K**). However, even if Hb binding can cause a substantial reduction in the enhancer activity, the net effect of including Hb binding sites in the enhancer is anyway shifted towards an increase in enhancer activity. The activity of short range repressors causes local histone deacetylation (22), which reduce DNA-accessibility in a region corresponding to roughly the size of one nucleosome (21). It has been known for a while that Hb can interact with the chromatin remodeler and de-acetylation complex NURD (34), thus already suggesting that Hb could act as a short-range repressor (33).

Overall, the simplest model to explain our observations assumes that Hb binding sites are able to increase enhancer activity by directly disfavoring nucleosome occupancy and thus increasing the enhancer accessibility. When these binding sites are occupied by Hb, expression is relatively reduced because of Hb activity as a short range repressor. However, the net balance between these effects still leans towards activation. Remarkably, we found the effect of the 3Hb sequence to be consistent when combined with two different groups of transcriptional activators. The fact that the effect of Hb seems to be independent from which activator targets the same enhancer, although limited to the cases of Bcd and Twi and Dl, is an important observation: it suggests that these processes are not driven by protein-protein interactions among TFs, a common mechanism of context dependent activity that was already proposed to explain Hb activity (30-32). Our observation of the action of Hb in Bcd driven enhancers allows to think the Hb-Bcd interaction as a simpler and more general effect due to Hb binding sites, instead of a direct interaction between the two proteins.

An alternative interpretation of our results could be given by assuming the presence of unintended binding sites in the 3Hb sequence. If present, these sites would have to be recognized by an ubiquitous activator like Zld or D-STAT. However, we didn’t find any strong binding sites in the 3Hb sequence for these or other factors that are known to be involved in the A-P or D-V axis segmentation (**Supplementary Fig. 3**). Therefore, one would have to postulate the existence of an additional TF recognizing a motif in the 3Hb sequence and synergistically increasing the effect of other activators, while not being able to drive any expression by itself. To explain our observations, this factor should also be homogenously expressed in the entire embryo in a constant amount during the blastoderm stage of development, since the 3Hb sequence does not influence enhancer dynamics but only expression levels. In addition, if sites for a general activator are present, one would expect them to act independently from binding sites orientation, which is not the case in our dataset.

In summary, we believe that an interpretation of these results based on a direct effect of Hb binding sites on nucleosome occupancy offers a simpler explanation for our observations. However, a direct proof of this mechanism would require further investigations based on different methods, in order to directly measure the impact of the 3Hb sequence as well as other sequences on both nucleosome occupancy and expression *in vivo*.

## MATERIALS and METHODS

### Generation of reporter constructs

Reporter constructs were cloned into an expression vector based on the pBDP backbone (a gift from Gerald Rubin; Addgene plasmid #17566) as described in (36) All enhancers were coupled to a strong synthetic core promoter (DSCP) driving the expression of the mNeon reporter (36). The sequence of all synthetic enhancers was generated by oligo annealing. In all constructs, except the two labeled “proximal”, a 73-bp linker separates the enhancer from the basal promoter. This sequence does not contain any predicted binding site for transcription factors of the segmentation network, as checked using PySite (available on *Github*: https://github.com/Reutern/PySite), a *Python* script we developed which uses positional weight matrices (PWMs) we determined a a previous work (40). A complete list of all sequences is provided in **Supplementary Information**. All vectors were integrated in the same attP2 docking site using PhiC31 integrase(51). Homozygous fly stocks were generated by crossing a single male with a single homozygous virgin female, and the insertion of the correct construct was verified by single-fly PCR of both parents and sequencing of the PCR products.

### Live Imaging

Enhancer-mNeonRep embryos were collected, dechorionated in 50% bleach and mounted between a semipermeable membrane and a microscope cover glass, immersed in halocarbon oil (Sigma). Imaging was performed at 24±1°C on a Zeiss LSM710 confocal microscope using a 40x 1.2NA water immersion objective. Pixel size was set to 1.1 µm. Two tiled stacks, each consisting of 3 images separated by 7.5µm in z, were acquired at each time point. The resulting field of view of 250 µm x 580 µm allowed us to image an entire embryo in a single movie with a time resolution of 60 s per z-stack.

### Image and data Analysis

Image and data analysis for the mNeon reporter has been described in detail in (36) and are briefly outlined here.

Confocal stacks of embryos were processed using the *Definiens XD 2*.*0* software package (Munich, Germany) to detect the cortical region of the embryo using the strong autofluorescence signal arising from their vitelline membrane (region delimited by the red lines in **Fig. 1B**). The mNeon green signal has been measured in this region in bins corresponding to 2% egg length. An average background autofluorescence profile, rescaled to account for autofluorescence photobleaching, was subtracted from the raw data.

Temporal registration, with a precision of ±1min, was achieved by detecting, in DIC images, the moment at which membranes reappear in the middle of the embryo after mitotic division following nc13.

Since mNeon fluorescence reflects protein level, which is an indirect readout of transcription, we reconstructed mRNA concentration from the time course of protein concentrations. To do so, we applied an image and data-analysis pipeline that we recently described in (36). Briefly, the analysis relies on a model of expression and maturation of the mNeonGreen reporter based on a set of linear ordinary differential equations (ODE). The model explicitly takes into account the rate of protein maturation and the degradation rates of both protein and mRNA, which have been previously characterized. Using linear inversion and a regularized non-negative least square algorithm, the ODE model then allows us to derive the mRNA concentration, the rate of instantaneous mRNA production, as well as the cumulative or total mRNA production from the reporter protein fluorescence for a given time window.

### Determination of DNA-histone binding free energy

We measured DNA-histone binding free energies with a fluorescence anisotropy assay we recently developed (42). Briefly, a robotic system was used to obtain competitive nucleosome formation of histones with a fluorescently labeled reference DNA sequence or a non-fluorescent competitor DNA sequence in a microwell plate. A modified epifluorescence microscope was used to measure fluorescence anisotropy in each well and derive the fraction of bound vs unbound DNA. Full titration curves were obtained by varying the concentration of the competitor sequence in different wells.

## ACKNOWLEDGMENTS

We dedicate this publication to the memory of Prof. Ulrike Gaul who passed away after a long illness during the completion of this work. The authors want to thank Prof. Nicolas Gompel, Prof. Don Lamb and Prof. David Arnosti for fruitful discussions about the project.

## AUTHORS CONTRIBUTIONS

SC, CJ, UU and UG developed the project; SC, UU and CJ designed the experiments; SC and MH generated fly stocks and performed in-vivo imaging of Drosophila embryos; MS performed nucleosome binding energy experiments; SC performed data analysis; SC and CJ wrote the manuscript.

## Supporting Information Captions

**Supplementary Figure 1:** 2Twi-2Dl enhancer with the original flanking sequence used in (21). The predicted binding sites for TFs of the segmentation network and the nucleosome occupancy for the 2Twi-2Dl enhancer. The heatmap in the first panel **(A)** represents binding sites strength at each position along the sequence. Two different color schemes are used to represent binding sites in the forward or reverse strands and thus the binding sites orientation. The lower panel **(B)** shows the predicted nucleosome occupancy over a larger region that covers the enhancer as well as the surrounding sequence, including the promoter, based on in vitro nucleosome sequence preference (17). The positions of the enhancer, promoter and TSS are highlighted with different shades of blue. Multiple strong binding sites for the segmentation TF Hb, highlighted with a red arrow in the figure, are present in the plasmid backbone, just upstream of the 2Twi-2Dl enhancer.

**Supplementary Figure 2:** 2Twi-2Dl enhancer with the neutral flanking sequence derived from (41). The predicted binding sites for TFs of the segmentation network and the nucleosome occupancy for the 2Twi-2Dl enhancer. The heatmap in the first panel **(A)** represents binding sites strength at each position along the sequence. Two different color schemes are used to represent binding sites in the forward or reverse strands and thus the binding sites orientation. The lower panel **(B)** shows the predicted nucleosome occupancy over a larger region that covers the enhancer as well as the surrounding sequence, including the promoter, based on in vitro nucleosome sequence preference (17). The positions of the enhancer, promoter and TSS are highlighted with different shades of blue.

**Supplementary Figure 3:** 3Hb-2Twi-2Dl enhancer with the neutral flanking sequence. The predicted binding sites for TFs of the segmentation network and the nucleosome occupancy for the 3Hb-2Twi-2Dl enhancer. The heatmap in the first panel **(A)** represents binding sites strength at each position along the sequence. Two different color schemes are used to represent binding sites in the forward or reverse strands and thus the binding sites orientation. The lower panel **(B)** shows the predicted nucleosome occupancy over a larger region that covers the enhancer as well as the surrounding sequence, including the promoter, based on in vitro nucleosome sequence preference (17). The positions of the enhancer, promoter and TSS are highlighted with different shades of blue.

**Supplementary Figure 4:** Effect of the 3Hb sequence on the shape of the expression profile of Bcd driven enhancers. **(A)** Fold change of the cumulative mRNA production driven by the Bcd driven synthetic enhancers compared to the Bcd3 enhancer. The fold change has been computed at each position and all time points and averaged. The homogeneity of the fold change throughout the AP axis, with the exception of the 3Hb-70-Bcd3 enhancer, proves that the 3Hb sequence rescales the expression pattern without altering its shape. **(B)** Normalized expression pattern of cumulative mRNA production driven by the Bcd driven synthetic enhancers. Expression for each enhancer has been normalized to its maximum at each time point.

**Supplementary Figure 5:** Effect of the 3Hb sequence at different distances from the promoter. **(A)** The cumulative mRNA production driven by the Bcd3-proximal positioned just upstream the DSCP promoter. **(B)** Cumulative mRNA production driven by the 3Hb-150-Bcd3-proximal positioned just upstream the DSCP promoter. (**C**) The 3Hb sequence at 150bp from 3Bcd binding sites induces a similar increase in expression level when the enhancer is positioned just upstream the promoter or further away.

**Supplementary Figure 6:** correlation between cumulative mRNA production driven by the 2Twi-2Dl, 3Hb-2Twi-2Dl and 3HbWeak-2Tw-2Dl enhancers and the in-vitro nucleosome binding energy of their enhancer sequence measured with a fluorescence anisotropy based assay. The cumulative mRNA production for the 2Twi-2Dl, 3Hb-2Twi-2Dl and 3HbWeak-2Tw-2Dl enhancers refers to the expression at the end of nc14 in the embryo posterior (55-75%AP), where Hb is not present. The error bars in the binding energy measurements represent a standard error of the mean over, on average, three replicates.

