## Supplemental figures and tables for "Context dependent activation and repression of enhancers by Hunchback binding sites in *Drosophila* embryo"

Stefano Ceolin et al.

**2Twi-2Dl Enhancer – Original Flanking**

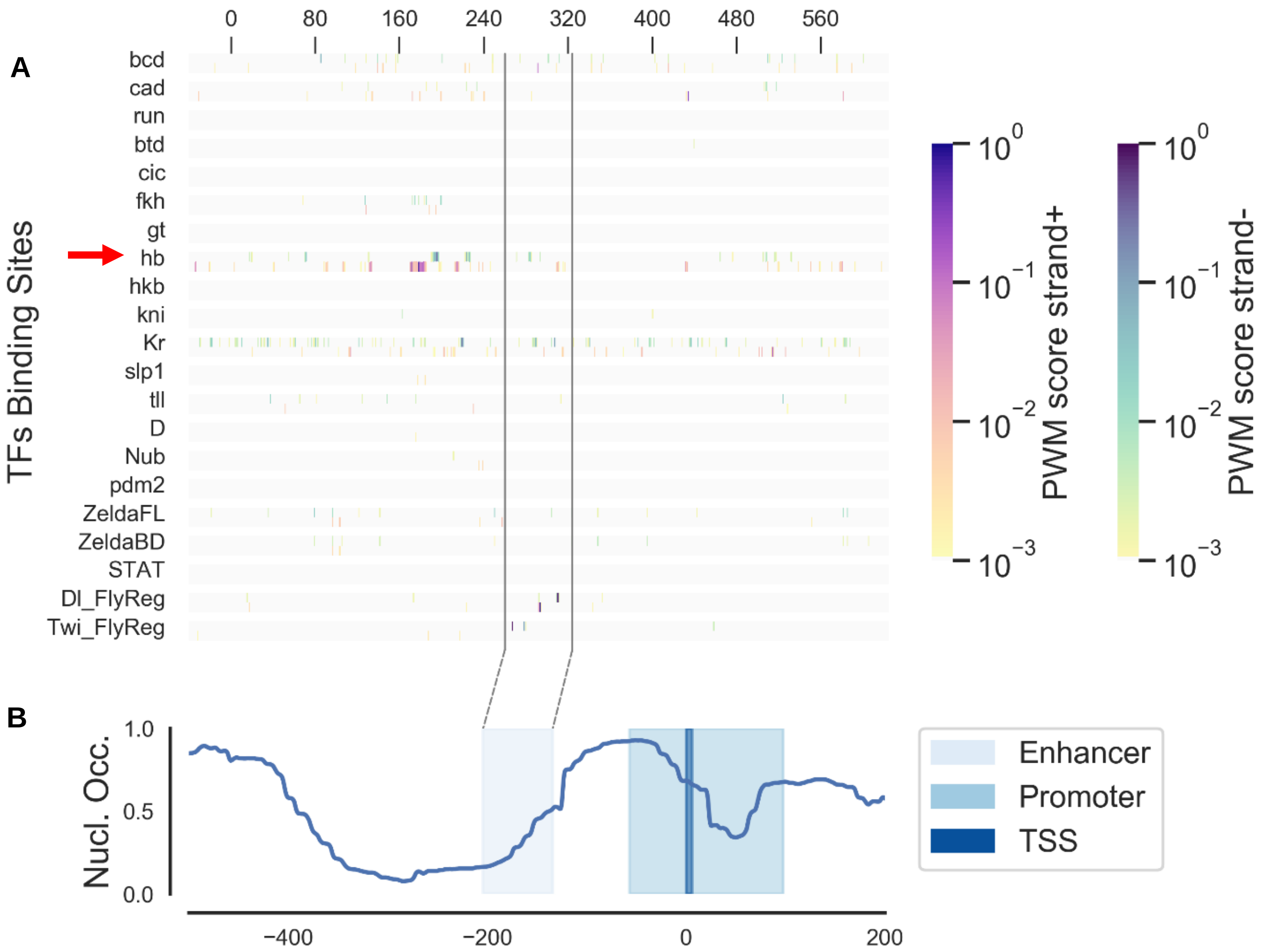


**Supplementary Figure 1:** 2Twi-2Dl enhancer with the original flanking sequence used in [1]. The predicted binding sites for TFs of the segmentation network and the nucleosome occupancy for the 2Twi-2Dl enhancer. The heatmap in the first panel **(A)** represents binding sites strength at each position along the sequence. Two different color schemes are used to represent binding sites in the forward or reverse strands and thus the binding sites orientation. The lower panel **(B)** shows the predicted nucleosome occupancy over a larger region that covers the enhancer as well as the surrounding sequence, including the promoter, based on in vitro nucleosome sequence preference [2]. The positions of the enhancer, promoter and TSS are highlighted with different shades of blue. Multiple strong binding sites for the segmentation TF Hb, highlighted with a red arrow in the figure, are present in the plasmid backbone, just upstream of the 2Twi-2Dl enhancer.

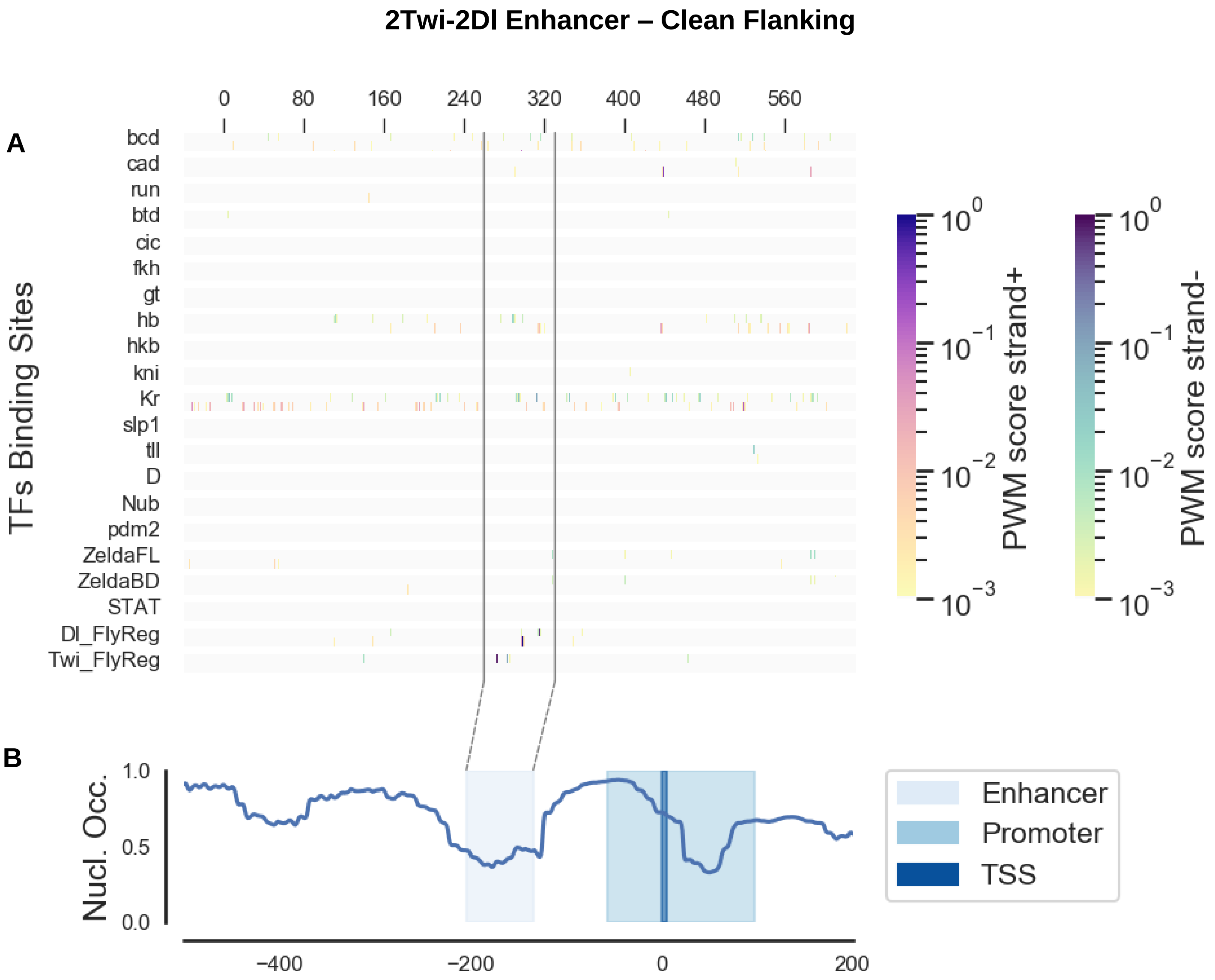


**Supplementary Figure 2:** 2Twi-2Dl enhancer with the neutral flanking sequence derived from [3]. The predicted binding sites for TFs of the segmentation network and the nucleosome occupancy for the 2Twi-2Dl enhancer. The heatmap in the first panel **(A)** represents binding sites strength at each position along the sequence. Two different color schemes are used to represent binding sites in the forward or reverse strands and thus the binding sites orientation. The lower panel **(B)** shows the predicted nucleosome occupancy over a larger region that covers the enhancer as well as the surrounding sequence, including the promoter, based on in vitro nucleosome sequence preference [2]. The positions of the enhancer, promoter and TSS are highlighted with different shades of blue.

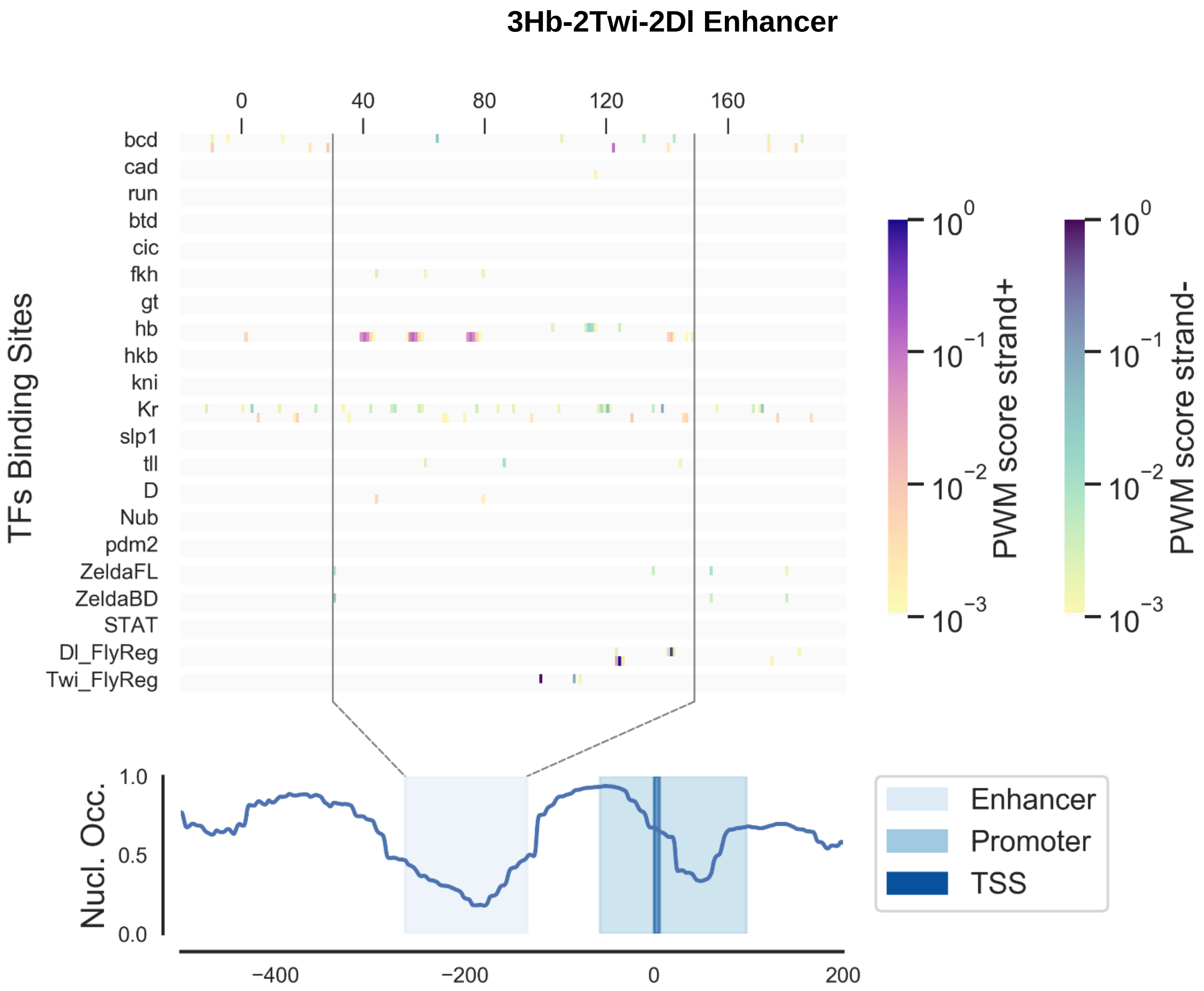


**Supplementary Figure 3:** 3Hb-2Twi-2Dl enhancer with the neutral flanking sequence. The predicted binding sites for TFs of the segmentation network and the nucleosome occupancy for the 3Hb-2Twi-2Dl enhancer. The heatmap in the first panel **(A)** represents binding sites strength at each position along the sequence. Two different color schemes are used to represent binding sites in the forward or reverse strands and thus the binding sites orientation. The lower panel **(B)** shows the predicted nucleosome occupancy over a larger region that covers the enhancer as well as the surrounding sequence, including the promoter, based on in vitro nucleosome sequence preference [2]. The positions of the enhancer, promoter and TSS are highlighted with different shades of blue.

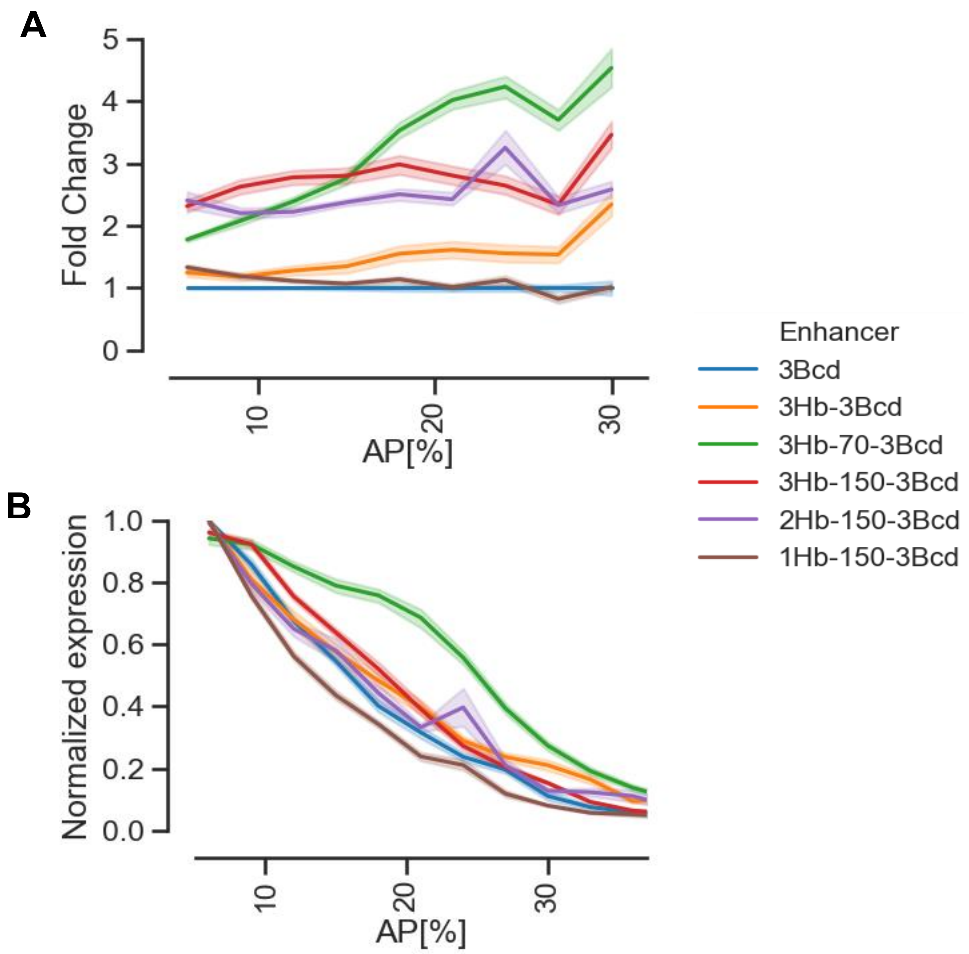


**Supplementary Figure 4:** Effect of the 3Hb sequence on the shape of the expression profile of Bcd driven enhancers. **(A)** Fold change of the cumulative mRNA production driven by the Bcd driven synthetic enhancers compared to the Bcd3 enhancer. The fold change has been computed at each position and all time points and averaged. The homogeneity of the fold change throughout the AP axis, with the exception of the 3Hb-70-Bcd3 enhancer, proves that the 3Hb sequence rescales the expression pattern without altering its shape. **(B)** Normalized expression pattern of cumulative mRNA production driven by the Bcd driven synthetic enhancers. Expression for each enhancer has been normalized to its maximum at each time point.

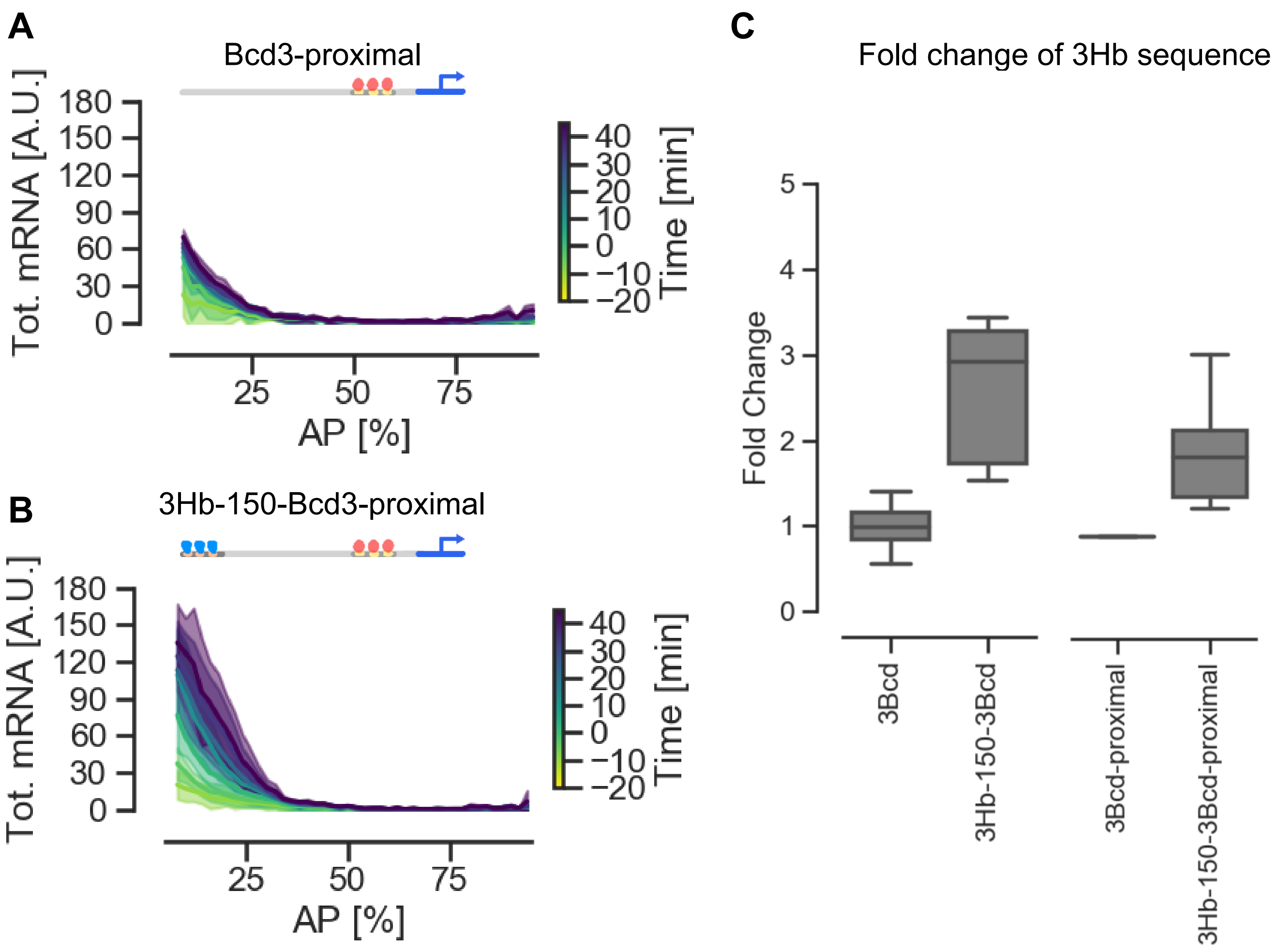


**Supplementary Figure 5:** Effect of the 3Hb sequence at different distances from the promoter. **(A)** The cumulative mRNA production driven by the Bcd3-proximal positioned just upstream the DSCP promoter. **(B)** Cumulative mRNA production driven by the 3Hb-150-Bcd3-proximal positioned just upstream the DSCP promoter. (**C**) The 3Hb sequence at 150bp from 3Bcd binding sites induces a similar increase in expression level when the enhancer is positioned just upstream the promoter or further away.

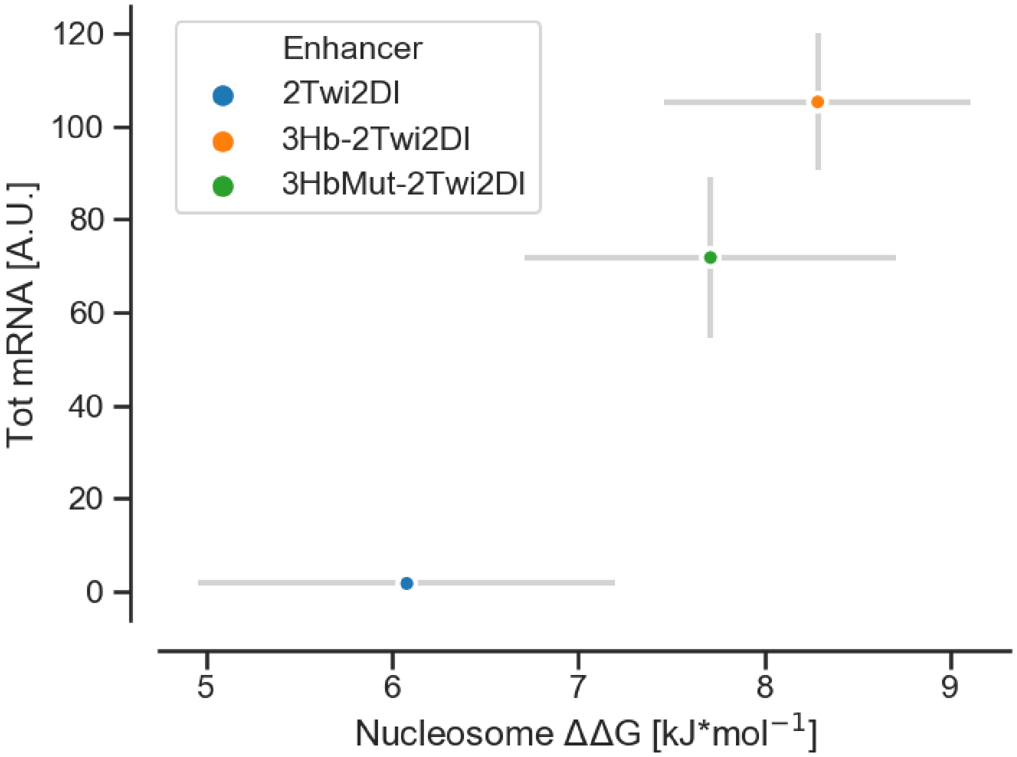


**Supplementary Figure 6:** correlation between cumulative mRNA production driven by the 2Twi-2Dl, 3Hb-2Twi-2Dl and 3HbWeak-2Tw-2Dl enhancers and the in-vitro nucleosome binding energy of their enhancer sequence measured with a fluorescence anisotropy based assay. The cumulative mRNA production for the 2Twi-2Dl, 3Hb-2Twi-2Dl and 3HbWeak-2Tw-2Dl enhancers refers to the expression at the end of nc14 in the embryo posterior (55-75%AP), where Hb is not present. The error bars in the binding energy measurements represent a standard error of the mean over, on average, three replicates.

### DNA Sequences

**Enhancers**

>2Twi-2Dl

CTCGCATATGTTGAGCATATGTTTTGGGGGATTTTCCCAAATCGAGGGAAAACC
CAA

>3Hb-2Twi-2Dl

AGGTACTCAAAAAACTATGGACTCAAAAAACTAATCGATACTCAAAAAACTATGA
GTCGAACTCGCATATGTTGAGCATATGTTTTGGGGGATTTTCCCAAATCGAGGG
AAAACCCAAACTAGT

>3Hb-70-2Twi-2Dl

AGGTACTCAAAAAACTATGGACTCAAAAAACTAATCGATACTCAAAAAACTATGA
GTCGAAACCGCTTTAGTCCCGCCAGCCAGCCACCAGTGGTGCAGCCCACCTCC
TCGGCCCGCCAGTCGCCCATCGCTCGCATATGTTGAGCATATGTTTTGGGGGAT
TTTCCCAAATCGAGGGAAAACCCAA

>3Hb-150-2twi-2Dl

AGGTACTCAAAAAACTATGGACTCAAAAAACTAATCGATACTCAAAAAACTATGA
GTCGAAACCGCTTTAGTCCCGCCAGCCAGCCACCAGTGGTGCAGCCCACCTCC
TCGGCCCGCCAGTCGCCCATCGATGTCTGCCTGGAGGAGGATGTTCACTCCGT
GCACAGCCATCAGTCGTCCGCAAGCCTCCTGCATCCCATTGCCATCCGACTCG
CATATGTTGAGCATATGTTTTGGGGGATTTTCCCAAATCGAGGGAAAACCCAA

>3Hb-250-2Twi-2Dl

AGGTACTCAAAAAACTATGGACTCAAAAAACTAATCGATACTCAAAAAACTATGA
GTCGAAACCGCTTTAGTCCCGCCAGCCAGCCACCAGTGGTGCAGCCCACCTCC
TCGGCCCGCCAGTCGCCCATCGATGTCTGCCTGGAGGAGGATGTTCACTCCGT
GCACAGCCATCAGTCGTCCGCAAGCCTCCTGCATCCCATTGCCATCCGAGCCA
CGCCAACCACTCCGACTAGCAGCAGCCCGCTGAGTTTTGCGGCCAAGATGCAG
AGCTTGTCGCCCGTTTCGGTTTGCTCCATTGGCGGCGAAACCACTCGCATATGT
TGAGCATATGTTTTGGGGGATTTTCCCAAATCGAGGGAAAACCCAA

>3HbWeak-2Twi2Dl

AGGTACTCAATAAACTATGGACTCAATAAACTAATCGATACTCAATAAACTATGA
GTCGAACTCGCATATGTTGAGCATATGTTTTGGGGGATTTTCCCAAATCGAGGG
AAAACCCAA

>3HbRev-2Twi-2Dl

TTCGACTCATAGTTTTTTGAGTATCGATTAGTTTTTTGAGTCCATAGTTTTTTGAG
TACCTCTCGCATATGTTGAGCATATGTTTTGGGGGATTTTCCCAAATCGAGGGAA
AACCCAA

>3Zld-2twi-2Dl

CGGAAGTTCAGGTATTGCTATTCAGGTAGAGGCCGTACGTGCAGGTAACCTTGC
GTCTCGCATATGTTGAGCATATGTTTTGGGGGATTTTCCCAAATCGAGGGAAAA
CCCAA

>Bcd3

AGGTTCTAATCCCGGTCTAATCCCTCGAGTCTAATCCCATGAGTCGAC

>Hb3-Bcd3

AGCTTAGGTACTCAAAAAACTATGGACTCAAAAAACTAATCGATACTCAAAAAAC
TATGAGTCGAAGGTTCTAATCCCGGTCTAATCCCTCGAGTCTAATCCCATGAGT
CGACG

>3Hb-70-3Bcd

AGGTACTCAAAAAACTATGGACTCAAAAAACTAATCGATACTCAAAAAACTATGA
GTCGAAACCGCTTTAGTCCCGCCAGCCAGCCACCAGTGGTGCAGCCCACCTCC
TCGGCCCGCCAGTCGCCCATCGAGGTTCTAATCCCGGTCTAATCCCTCGAGTCT
AATCCCATGAGTCGACG

>3Hb-150-3Bcd

AGGTACTCAAAAAACTATGGACTCAAAAAACTAATCGATACTCAAAAAACTATGA
GTCGAAACCGCTTTAGTCCCGCCAGCCAGCCACCAGTGGTGCAGCCCACCTCC
TCGGCCCGCCAGTCGCCCATCGATGTCTGCCTGGAGGAGGATGTTCACTCCGT
GCACAGCCATCAGTCGTCCGCAAGCCTCCTGCATCCCATTGCCATCCGAAGGTT
CTAATCCCGGTCTAATCCCTCGAGTCTAATCCCATGAGTCGACG

>2Hb-150-3Bcd

ACTCAAAAAACTAATCGATACTCAAAAAACTATGAGTCGAAACCGCTTTAGTCCC
GCCAGCCAGCCACCAGTGGTGCAGCCCACCTCCTCGGCCCGCCAGTCGCCCA
TCGATGTCTGCCTGGAGGAGGATGTTCACTCCGTGCACAGCCATCAGTCGTCC
GCAAGCCTCCTGCATCCCATTGCCATCCGAAGGTTCTAATCCCGGTCTAATCCC
TCGAGTCTAATCCCATGAGTCGACG

>1Hb-150-3Bcd

TACTCAAAAAACTATGAGTCGAAACCGCTTTAGTCCCGCCAGCCAGCCACCAGT
GGTGCAGCCCACCTCCTCGGCCCGCCAGTCGCCCATCGATGTCTGCCTGGAG
GAGGATGTTCACTCCGTGCACAGCCATCAGTCGTCCGCAAGCCTCCTGCATCC
CATTGCCATCCGAAGGTTCTAATCCCGGTCTAATCCCTCGAGTCTAATCCCATG
AGTCGACG

>3HbW-150-3Bcd

AGGTACTCAcAAAACTATGGACTCAcAAAACTAATCGATACTCAcAAAACTATGAG
TCGAAACCGCTTTAGTCCCGCCAGCCAGCCACCAGTGGTGCAGCCCACCTCCT
CGGCCCGCCAGTCGCCCATCGATGTCTGCCTGGAGGAGGATGTTCACTCCGTG
CACAGCCATCAGTCGTCCGCAAGCCTCCTGCATCCCATTGCCATCCGAAGGTTC
TAATCCCGGTCTAATCCCTCGAGTCTAATCCCATGAGTCGACG

>3HbRev-150-3Bcd

TCGACTCATAGTTTTTTGAGTATCGATTAGTTTTTTGAGTCCATAGTTTTTTGAGT
ACCTAACCGCTTTAGTCCCGCCAGCCAGCCACCAGTGGTGCAGCCCACCTCCT
CGGCCCGCCAGTCGCCCATCGATGTCTGCCTGGAGGAGGATGTTCACTCCGTG
CACAGCCATCAGTCGTCCGCAAGCCTCCTGCATCCCATTGCCATCCGAAGGTTC
TAATCCCGGTCTAATCCCTCGAGTCTAATCCCATGAGTCGACG

>3HbRev-3Bcd

TCGACTCATAGTTTTTTGAGTATCGATTAGTTTTTTGAGTCCATAGTTTTTTGAGT
ACCTAGGTTCTAATCCCGGTCTAATCCCTCGAGTCTAATCCCATGAGTCGACG

**Linkers**

Enhancer-promoter linker sequence:

>Linker

AGGTTCCAGTGTGGTGGAATTCTGCAGATATCCAGCACAGTGGCGGCCGCTCG
AGTCTAGAGGGCCCTTCGAA

Spacer sequence, inserted upstream all enhancer sequences to create a neutral background around enhancers:

>Neutral_Spacer

GAATTCGGCCGGCCCACCAAACCGCTTTAGTCCCGCCAGCCAGCCACCAGTGG
TGCAGCCCACCTCCTCGGCCCGCCAGTCGCCCATCGATGTCTGCCTGGAGGAG
GATGTTCACTCCGTGCACAGCCATCAGTCGTCCGCAAGCCTCCTGCATCCCATT
GCCATCCGAGCCACGCCAACCACTCCGACTAGCAGCAGCCCGCTGAGTTTTGC
GGCCAAGATGCAGAGCTTGTCGCCCGTTTCGGTTTGCTCCATTGGCGGCGAAA
CCACCAGCGTTGTACCAGTGCATCCTCCCACCGTTTCCGCTCAAGAAGGACCCA
TGGATCTGAGCATGAAGACCTCGCGGAGCTCCGT

Flanking sequence from the reporter plasmid used in [1], inserted upstream the 2Twi-2Dl enhancer in the Original-2Twi-2Dl construct.:

>Original_Flanking

AAGCTTGCACAGCACTTTGTGTTTAATTGATGGCGTAAACCGCTTGGAGCTTCG
TCACGAAACCGCTGACAAAATGCAACTGAAGGCGGACATTGACGCTACGTAACG
CTACAAACGGTGGCGAAAGAGATAGCGGACGCAGCGGCGAAAGAGACGGCGA
TATTTCTGTGGACAGAGAAGGAGGCAAACAGCGCTGACTTTGAGTGGAATGTCA
TTTTGAGTGAGAGGTAATCGAAAGAACCTGGTACATCAAATACCCTTGGATCGA
AGTAAATTTAAAACTGATCAGATAAGTTCAATGATATCCAGTGCAGTAAAAAATAA
AAAAAAAATATGTTTTTTTAAATCTACATTCTCCAAAAAAGGGTTTTATTAACTTAC
ATACATACTAGAATTCGGTACCGAG
